# Thermodynamic constraints are sufficient for the emergence of flux sensors in metabolism

**DOI:** 10.1101/2020.09.04.283010

**Authors:** Christian Karl Euler, Radhakrishnan Mahadevan

## Abstract

Metabolism is a precisely coordinated phenomenon, the apparent goal of which is to balance fluxes to maintain robust growth. However, coordinating fluxes requires information about *rates*, which is not obviously reconcilable with known regulatory mechanisms in which *concentrations* are sensed through metabolite binding. While flux sensor examples have been characterized, the fundamental principles underlying the phenomenon in general are not well understood. Specifically, the questions of which fluxes can be sensed, and the mechanism by which they are remain open. We address this by showing that the concentrations of substrates of thermodynamically constrained reactions reflect upstream flux and therefore carry information about rates which can be propagated through regulatory interactions to control other fluxes in the network. Using fluxomic, metabolomic, and thermodynamic data in *E coli*, we show that the concentrations of a few metabolites in central carbon metabolism reflect their producing fluxes and demonstrate that they can transmit information about these rates because of their positions in the network and their roles as effectors.

## Introduction

All cells must be able to take in material from their environments and coordinate its reconstitution into the necessary components in order to persist (***Chubukov et al., 2014***). Growth rate is the primary metric by which the ability of cells to perform this task is assessed: metabolisms which enable higher and more robust *rates* for a given niche have been selected for over evolutionary timescales. This poses a logical challenge, because there is no obvious way for cells to directly measure rates, and yet control of fluxes appears to be a goal – perhaps *the* goal – of metabolism (***Feist and Palsson, 2010***). The principle of observability would suggest that control of rates would be difficult, if not impossible, to achieve without at least some of those rates being observable (***Kalman, 1960***). However, all known regulatory interactions in living systems are based on the concentration-driven binding of small molecules to enzymes, transcription factors, etc. to modulate their activity. That is, signals directly available to biological control systems appear to be amounts, not rates. These systems control metabolic fluxes through direct changes to protein activity and/or changes in enzyme levels. In this sense, metabolites are informational signals, and all environmental information must be encoded in their amounts (***Wegner et al., 2015***). Understanding how cells decode these signals to both generate information, and make decisions about rates to effectively partition resources is essential to understanding metabolism. How, then does the effective control of rates arise in metabolic networks which can only sense internal amounts?

A metabolic signal which grows with a flux or fluxes could easily be used as the input to a regulatory system to achieve the goals of flux sensing and regulation (***Litsios et al., 2018***). Indeed, recent experimental work has uncovered that the concentrations of the small molecules fructose-1,6-bisphosphate (FBP) and cyclic AMP (cAMP) are functions of fluxes in *E coli* and as such are useful as signals of these rates. FBP is a signal of total carbon flux through upper glycolysis, regardless of its source, and allosterically regulates several enzymes involved in carbon utilization. These regulatory interactions coordinate fluxes in upper and lower glycolysis and enable hierarchical carbon source use (***Kochanowski et al., 2013***; ***Okano et al., 2020***). Similarly, cAMP concentration decreases linearly with increasing growth rate along the so-called “C-line”. The cAMP-crp system coordinates the expression of biosynthetic genes with growth rate through transcriptional regulation (***You et al., 2013***). These two examples establish flux-sensing as an integral part of the decision-making process for coordinating both carbon uptake and gene expression with growth rate in *E coli* (***Okano et al., 2020***). FBP also plays roles in signalling glycolytic flux to signal transduction systems associated with proliferation in *S cerevisiae* (***Peeters et al., 2017***).

While these molecules are to date the only confirmed flux signals, it is well known that some metabolic processes occur as functions of a flux or fluxes. For example, overflow metabolism in both *S cerevisiae* (***Huberts et al., 2012***) and *E coli* switches on as a function of total glycolytic flux, not carbon source or amount. In *E coli*, cAMP coordinates the concomitant proteomic shifts in the transition to aerobic fermentation, whether these shifts are causative or not (***Basan et al., 2015***). The molecular origins of overflow metabolism are still unsettled, but these observations suggest it is governed by *rates*, not *amounts*. Other recent work suggests that perhaps flux sensing in the case of the shift to overflow metabolism arises due to constraints on the ability of metabolic networks as a whole to dissipate energy, i.e. that it emerges from systemic thermodynamic constraints on metabolism (***Niebel et al., 2019***). The ability for metabolisms to sense rates through concentrations is ostensibly a fundamental aspect of these networks, and perhaps essential to their stability and propagation, yet there is currently a dearth of theory about the origins of the phenomenon.

This poses a challenge to engineering efforts. In metabolic engineering, the control of flux distributions is required to optimize production of chemical targets. Some targets require precise splitting and recombination of fluxes to control the ratio of precursor supply, and such control could be achieved with a flux sensor-actuator (***Gold et al., 2015***). Beyond that, when engineers perturb metabolic processes there is potential for disrupting native, unknown flux sensing relationships with deleterious effect. Understanding where such relationships exist or could exist could help to assess their robustness to the large changes in fluxes that engineers often attempt in the course of overproducing target chemicals. Finally, real-time flux observations via fluorescence signals could find uses in medical research, for example in the examination of cancer and/or stem cell metabolisms under perturbation (***Peeters et al., 2017***). To this end, a synthetic biology tool which makes use of FBP as an input signal for a fluorescent output has been developed (***Monteiro et al., 2019***). This highlights the potential for making use of native metabolites which are sensors of fluxes as a way to control and/or observe metabolism for a variety of purposes.

In this work, we begin by assuming that flux sensing could be a relatively widespread phenomenon in metabolism. With this in mind, we consider the general conditions on reactions which could lead to feasible correlative relationships between fluxes and metabolites. We demonstrate that the concentrations of substrates of reactions for which rates are constrained can reflect their producing fluxes without the previously-suggested requirement of feedforward regulation (***Kochanowski et al., 2013***). The signals generated by these metabolites can be integrated by local small-molecule regulatory interactions to change downstream fluxes in response to changes in upstream fluxes. Then we make use of existing metabolomic and fluxomic data from *E coli* to show that citrate, malate, and some pentose phosphate pathway intermediates are very likely flux sensors, that phosphoenolpyruvate may be as well, and that glycerol-3-phosphate satisfies the structural criteria for flux-sensing. All of these are involved in the flux signal generation and interpretation motifs we propose. Further, we suggest that the flux-sensing functions of malate, citrate, and phosphoenolpyruvate are dependent on environmental conditions based on the sensitivity of the small-molecule regulatory interactions in which they are involved, and differential expression of downstream regulatory targets. The concentrations of the flux-sensing candidates we identify appear to be sufficient to predict fluxes in the major pathways in *E coli* central carbon metabolism and coordinate these fluxes via flux coupling.

## Results

### Enzyme Capacity Defines Substrate Sensitivity to Changes in Flux

We propose three criteria for metabolites to be considered flux sensors. First, they must have concentrations which increase monotonically as a given flux or fluxes increase (***Litsios et al., 2018***). In general this is not guaranteed for an arbitrary metabolite in a complex network because concentrations are dependent on local topology and the history of the system. Thus, flux-concentration relationships should arise due to the structure of the network. Second, an ideal flux sensor should have a linear or near-linear response which reasonably amplifies flux signals without causing or requiring extreme conditions. Finally, flux signalling metabolites should have concentrations which are constrained to be high under some conditions, but not others. Heterogeneity in concentration across different conditions means that a metabolite could be a useful internal signal of the fluxes associated with those conditions, and can therefore be used to make decisions which distinguish between them (Fig 1A).

**Figure 1.**
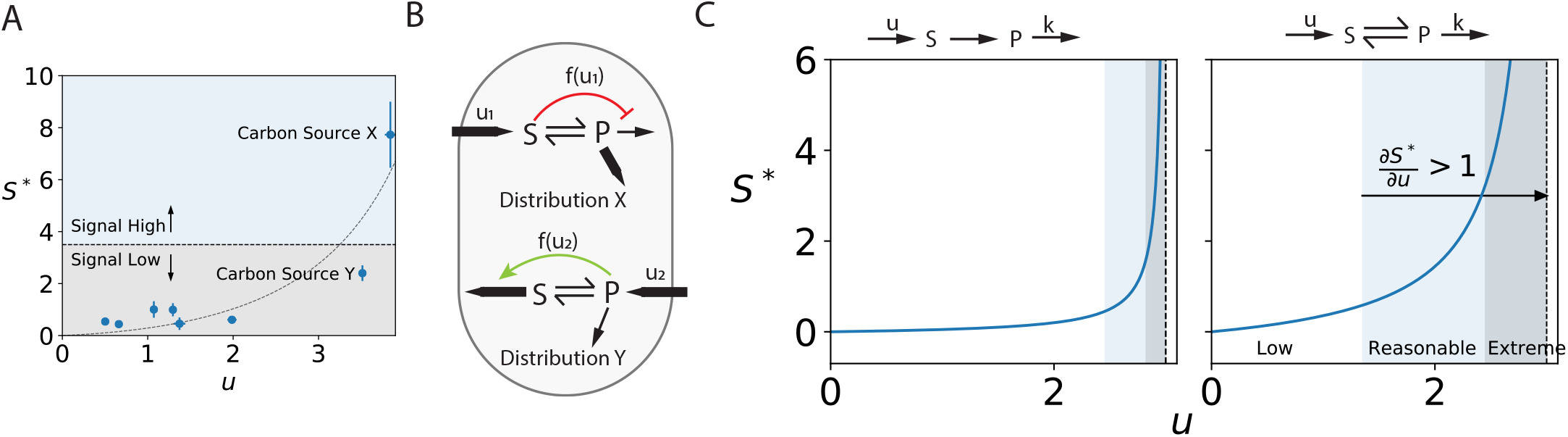
(A) Concentration-flux profile for a generic flux sensor. The steady-state concentration, *S*^∗^, increases monotonically as its producing flux *u* increases. Regulatory parameters, such as *K*_*I*_ values, define signal strength so that high and low fluxes can be distinguished (dashed line). For example, *S* could be the assimilation product of Carbon Source X, with *u* reflecting the uptake of this carbon source. On other carbon sources, *S*^∗^ is below the threshold, reflecting a low uptake rate. (B) Information carried by flux sensors can be fed forward into the network to effect changes in the downstream flux distribution which are consistent with use of the new carbon source. Red indicates inhibition, and green indicates activation. (C) Constraints on flux capacity and direction define the structural conditions in which a metabolite is a flux sensor. The sensitivity of *S*^∗^ to changes in 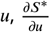, divides the *S* − *u* curve into three regions. In the low amplification region, coloured white, 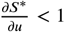. Signals are reasonably amplified when 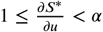 for some *α* > 1, here coloured blue. Extreme signal amplification, coloured grey, occurs for 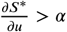, as *u* → *V*_*m*_ for both cases. Here, amplification of flux signals could cause instabilities in the network by forcing large concentrations of *S*.We arbitrarily chose 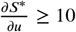 as our criteria for this region for comparison purposes. The reasonable amplification region is expanded for reactions which are favoured in the reverse direction (right panel) relative to those which are highly favoured in the forward direction (i.e. irreversible, left panel). *V*_*m*_ =3 and *K*_*s*_ = 0.1 for both simulations. Kinetics of the form in equation 1 were used in the left panel, while those of the form in equation 4 with *K*_*eq*_ = 0.01 and *K*_*p*_ =1 were used in the right panel.

To understand the kinetic constraints which could guarantee that a metabolite meets these criteria, we first consider the simple enzymatic transformation of a substrate *S* to product *P* in which *S* arrives at an arbitrary influx rate of *u* independent of the rate at which it is consumed and which may be the sum of many individual rates, and *P* has a turnover proportional to *k*. This is representative of one step in a linear metabolic pathway. Assuming simple Michaelis-Menten kinetics and only one consuming reaction, the mass balance for *S* is

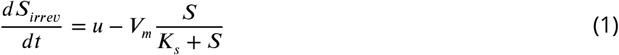

And the steady state concentration of *S*, 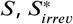, as a function of the influx *u* is given by:

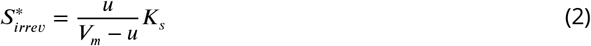

With non-negativity guaranteed for *u* ≥ 0. Biologically feasible solutions also require that *u* be sufficiently smaller than *V*_*m*_, otherwise the substrate will accumulate indefinitely (Fig 1C, grey region). In reality, all metabolite concentrations are bounded from above due to physical constraints, so indefinite accumulation is not physically realizable.

Equation 2 demonstrates that there is a relationship between the steady state concentration of the substrate and its influx over the range of possible influxes, so even this simple reaction scheme may meet the first criterion for flux sensing (Fig 1C). We can characterize the strength of this flux-concentration relationship by examining how 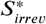 changes with changes to *u*:

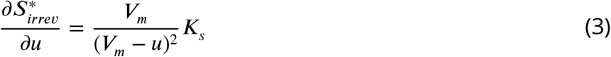

As in Fig 1, this sensitivity can be used to divide the *S* − *u* relationship into three regions. For low influxes, 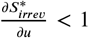 corresponding to low flux sensitivity. Here the relationship between influx and concentration is approximately linear, but any perturbations to upstream flux only weakly affect the concentration of *S*. When 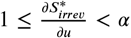 for some *α* > 1, changes to *u* are reasonably amplified in the concentration of *S* (blue region, Fig 1C). This is desirable for a flux sensor and occurs for 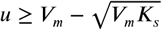. In the irreversible case, this region is a small fraction of the possible influxes and very small changes in *u* here could drive the system into the region of extreme signal amplification which occurs for 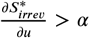 (grey region, Fig 1C). Here, the enzyme is nearly saturated and its capacity is therefore minimally dependent on substrate concentration. As a result, very large changes in the steady state substrate concentration may occur in response to small changes in influx. *V*_*m*_ can be adjusted via changes to enzyme expression to avoid unreasonable accumulation of *S* and the extreme sensitivity scenario. However, increases to *V*_*m*_ reduce the sensitivity of the substrate concentration to *u*, because as *V*_*m*_ increases the region of influxes for which 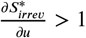 decreases.

Substrates could act as flux sensors in any of the three sensitivity regions, because regulatory interactions can be tuned to any particular concentration of *S*. However, local systems which typically operate in the 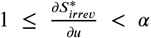 region meet our criteria for reasonable signal amplification, because they turn small changes in flux into larger changes in concentrations without causing excessive signal amplification or risking extreme accumulation of *S*. In practice, there is no obvious choice of value for *α*, but a value corresponding to *u* as some fraction of *V*_*m*_ could be taken. For example 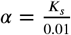 corresponds to *u* = 0.9*V*_*m*_ for the upper bound on reasonable signal amplification.

These results suggest that substrates consumed by individual irreversible reactions can yield reasonable flux-sensing scenarios for a narrow range of values of *K*_*S*_ and *V*_*m*_. However, in real metabolic networks metabolites are typically substrates for many reactions with different products. We can thus consider the situation in which *S* is consumed by many irreversible reactions. In this case, it will not be sensitive to *u* until all but one downstream enzyme is near saturation (Fig 2A). Up until this point, there will always be downstream capacity for increases to influx, so the existence of multiple consuming reactions increases the feasible range of 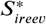 up to the sum of all of the capacities of the consuming reactions (i.e. up to *u* = Σ*V*_*m*.*i*_). This significantly expands the feasible range of influxes, but constrains the range over which the substrate concentration is sensitive to changes in influx by expanding the low sensitivity region. Substrates of branched pathways are therefore less likely to be useful as flux sensors (see Appendix 1). Thus we can conclude that in general flux sensing could emerge from kinetically constrained reactions, but that in this case it trades off with infeasible accumulation of substrate and is dependent on both expression of downstream enzymes and local network topology. As a result, kinetically constrained scenarios fail to fully meet even the first criterion for flux senors.

**Figure 2.**
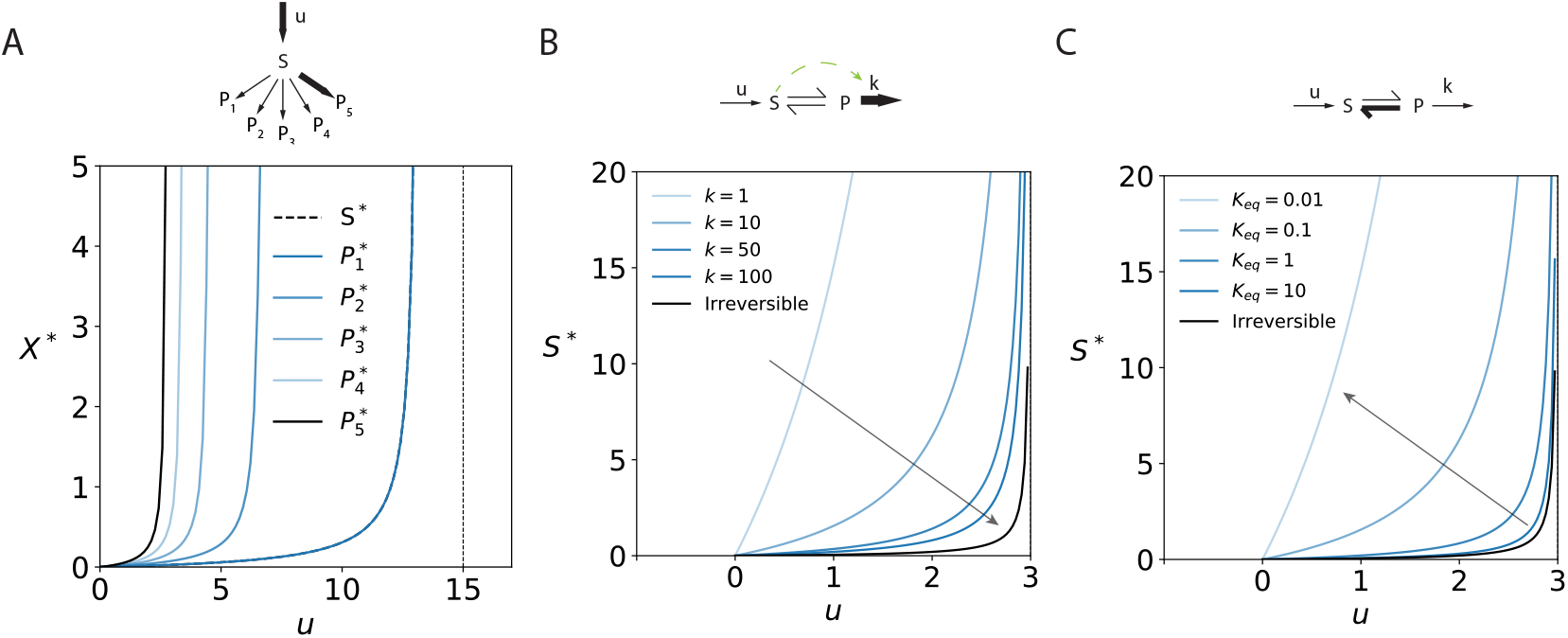
(A) Flux-concentration relationship for a branched, irreversible pathway. Downstream pathways saturate sequentially according to flux capacity (*V*_*P* 5_ > *V*_*P* 4_ > *V*_*P* 3_ > *V*_*P* 2_ > *V*_*P* 1_). Accumulation of *S* only occurs when *u* → Σ*V*_*m*_ (15 here), so substrates with multiple consuming pathways are less likely to behave as flux sensors. *K*_*s*_ = 0.1 for all consuming enzymes. (B) High downstream turnover (large *k*) can overcome thermodynamic bottlenecks. Arbitrary increases to *k* reduce the steady substrate concentration (*S*^∗^) for reversible reactions such that the *S*^∗^ profile approaches that of the irreversible case. Arrow indicates direction of increase to *k*. Increases to downstream turnover may be achieved through positive regulation of the consuming enzymes of *P*, as indicated with the dashed green arrow (***Kochanowski et al., 2013***). *V*_*m*_ = 3, *K*_*s*_ = 0.1, *K*_*p*_ = 1, *K*_*eq*_ = 0.1. (C) Degree of reversibility defines the range over which *S*^∗^ is a linear function of influx, *u*. For reactions highly favoured in the reverse direction (*K*_*eq*_ ≪ 1), the response of *S*^∗^ is steep (proportional to 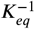) and approximately linear over the entire stable range. This is the ideal concentration-flux relationship for a flux-sensing metabolite as it amplifies the flux signal and provides a linear signal for regulation. Arrow indicates decreasing *K*_*eq*_. *V*_*m*_ = 3, *K*_*s*_ = 0.1, and *k* = 10.

In contrast, thermodynamic constraints caused by reactions which are unfavourable in the direction compatible with metabolic function cannot be avoided by changing enzyme expression. In general, the evolutionary innovation addressing thermodynamic bottlenecks is the coupling of reversible reactions to high-energy carrier molecules such as ATP and NADH to drive them in the desired direction. However, this strategy is ostensibly not a general solution, as evidenced by the persistence of reverse-favoured reactions in metabolic networks. Because these reactions require large mass action ratios to be driven in the desired direction, we suspected that they could yield stable flux-concentration relationships useful for control.

Therefore we now consider the reversible transformation of *S* to *P* with similar parameterization as before. The relevant mass balances for this scheme are

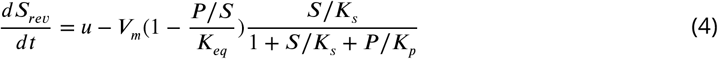

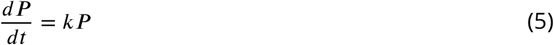

Where *K*_*eq*_ is the equilibrium constant for the reaction (***Noor et al., 2013***). At steady state, all net fluxes are equal to the influx, *u*, so the steady state substrate concentration, 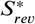, is

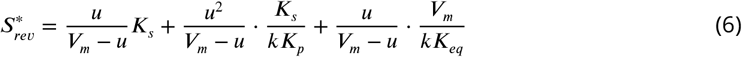

Once again, *u* < *V*_*m*_ is required to avoid indefinite accumulation of *S*, and *u* ≥ 0 for non-negativity. Equation 6 can be rearranged and scaled to the value for the equivalent irreversible case, 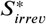, to highlight the behaviour of 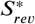 over *u*:

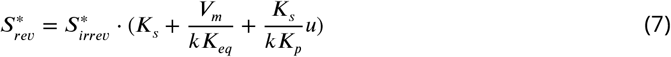

Since the first term in equation 6 is equivalent to equation 2 and the other two terms are guaranteed to be positive, the reversible steady state substrate concentration is guaranteed to be greater than or equal to that of the irreversible case, 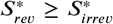 (Fig 2C). Equality only occurs when both *k* and *K*_*eq*_ are large, reflecting large downstream turnover and a favourable forward reaction. In practice, the final term in equation 6 dominates, so we can use this to understand exactly when 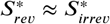, which is also the condition for 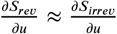:

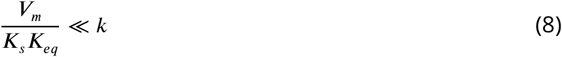

Thus, the downstream turnover rate must be many times larger than that of the reversible step in order for the reversible steady state concentration profile to approach that of the irreversible case (Fig 2B). This highlights the fact that downstream kinetic pull can help overcome thermodynamic constraints, reducing the strength of flux sensing relationships, but that the degree to which this is possible is constrained by the thermodynamic nature of the given reaction. As an example, for a reaction with *K*_*eq*_ ∼ 0.01 (e.g. aconitase), the downstream turnover rate would have to be much larger than 100x that of the reversible step for 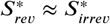.

Similarly to the irreversible case, we can examine the sensitivity of the steady state substrate concentration to changes in influx:

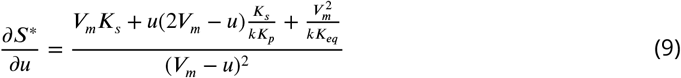

Equation 9 can be rearranged to the form:

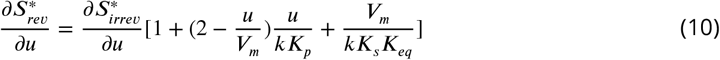

As in the irreversible case, equation 10 can be used to divide the *S* − *u* curve into three regions: linear, reasonable amplification, and extreme amplification (Fig 1B). Because 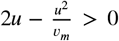 for 0 ≤ *u* < *V*_*m*_, the sensitivity is guaranteed to be larger in the reversible case than in the irreversible case for all *u* except in the condition given in equation 8. As a result, both the region for which the flux signal is reasonably amplified, and the region of extreme amplification are expanded. The fraction of the total sensitive region corresponding to extreme amplification is a function of 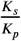 in this case, with low forward sensitivity yielding a larger fraction relative to the irreversible case. For reverse-favoured reactions, the extreme amplification region may make up a larger portion of the sensitive region for any choice of *α*, but this effect can be minimized if the enzyme is more sensitive to substrate than it is to product in the forward direction (see Appendix 1).

Both kinetic and thermodynamic constraints on flux facilitate linear flux-concentration relationships for *u* ≪ *V*_*m*_ which are defined by their sensitivities at *u* = 0. In the reversible case, the sensitivity is dependent on the degree to which the reverse reaction is favoured:

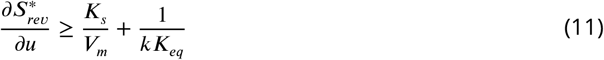

With the first term of equation 11 being equal to that of the irreversible case. This means that the more favourable the reverse reaction is, the stronger and more linear the relationship between perturbations to influx and the sensible concentration of the substrate is. Thus, thermodynamic constraints are not only yield amplified signals for a wider range of fluxes, but the degree to which they constrain forward flux also controls the linearity of the flux signals they produce (Fig 2C). Reducing *V*_*m*_ in the irreversible case is required to achieve the same effect as increasing favourability of the reverse reaction in the reversible case: in effect, reactions which are favoured in the reverse direction decouple flux sensing properties from kinetic capacity, eliminating the tradeoff that exists in the kinetically constrained case.

As in the irreversible case, we can again consider the situation in which *S* is consumed by many reactions, one of which is reversible. In the reversible case, all of the irreversible reactions are guaranteed to saturate before the reversible reaction because it will contribute to the accumulation of substrate through backflux up to 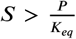. Thus in this situation the reversible reaction is guaranteed to constrain the steady state substrate concentration. A metabolite must only be consumed by a single reaction favourable in the reverse direction for it to satisfy the structural properties of a flux sensor that we have outlined here.

Our analysis reveals that metabolites which are involved in reactions for which flux capacity is kinetically constrained have potential to be flux sensors because their concentrations reflect the magnitude of an upstream flux/fluxes and they can amplify perturbations to these rates. However, the flux-sensing performance of these reactions is not guaranteed by network structure, but is condition-dependent. In contrast, thermodynamically constrained reactions yield stronger flux signals over a wider range of fluxes than those constrained by kinetics, and they do so as a function of the degree of constraint (i.e. *K*_*eq*_), rather than as a function of a condition-dependent parameter (i.e. *V*_*m*_). While it was previously claimed that feedforward activation is required for flux-sensing in the FBP case (***Kochanowski et al., 2013***), we have shown that it is sufficient for FBP to be the substrate of a reaction which is favourable in the reverse direction, fructose bisphosphate aldolase (FBA, *K*_*eq*_ ∼ 10^−4^ (***Flamholz et al., 2012***)) for it to be a signal of glycolytic flux.

If feedforward activation (FFA) is not necessarily a requirement for flux sensing, then an open question is what is the role of such regulatory interactions when they involve flux-sensing metabolites? Here, we argue that FFA – and other small molecule regulatory motifs – arise not because they are required to make metabolites sensors of flux, but rather that they serve to capture information encoded by metabolites which are flux sensors as a result of local constraints. In this way, small molecule regulatory (SMR) interactions could coordinate fluxes by regulating some fluxes as a function of others. This question and hypothesis are investigated next.

### Feedforward Structures Can Interpret Flux Signals

Flux signals emerging from local constraints must be interpreted elsewhere in the network in order to effect control action. SMR is the likely regulatory mode by which rate information is decoded at the metabolic level, because it involves direct modulation of fluxes through changes to enzyme activity (***Litsios et al., 2018***). It is also consistent with the FFA of pyruvate kinase by FBP in which a rate signal is used to to control other rates through SMR. Simple FFA has been explored in depth and its dynamics are well understood: it linearizes and stabilizes responses by increasing downstream turnover as a function of upstream flux (***Kochanowski et al., 2013***; ***Hovd and Bitmead, 2009***). Therefore, here we will explore two other feedforward SMR structures which would enable decision-making with flux signals as inputs: splits and recombines. In the former, one flux is distributed among two or more downstream fluxes (Fig 3A), while in the latter two or more fluxes combine at a node (Fig 3B). Both of these structures are types of branchpoints, but they differ in directionality.

**Figure 3.**
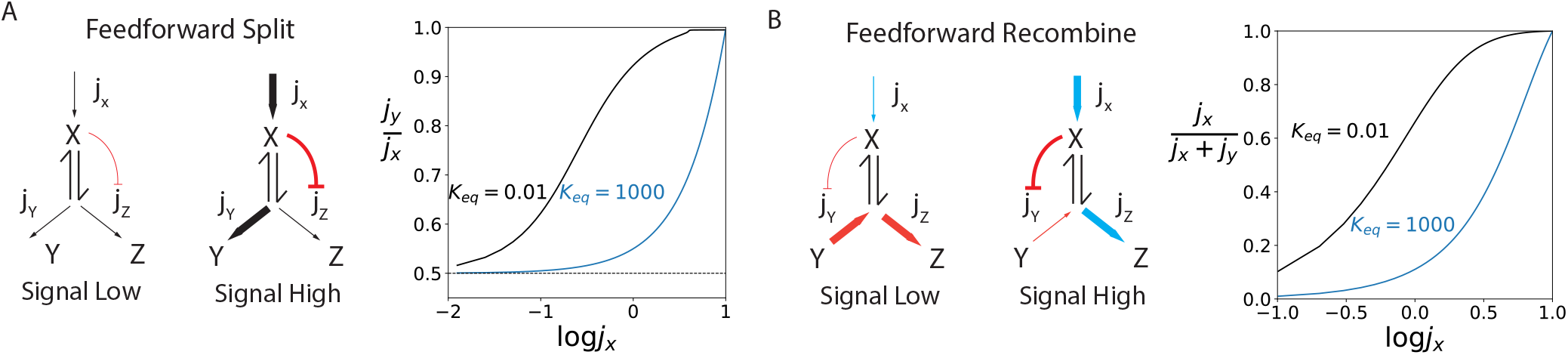
Small-molecule regulatory (SMR) motifs make use of thermodynamic constraints for flux partitioning. (A) In the feedforward split architecture, influx, *j*_*x*_, is split among two pathways, with one branch (*j*_*z*_ here) being regulated by the flux-sensing metabolite *X*. The proportion of flux through the unregulated pathway, 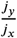 is always a function of total influx when is the substrate of a thermodynamically constrained reaction (black line), but only at very large influx values when it is not (blue lines). Since the two paths share a common source, there is a baseline ratio of flux to either which can only be changed through changes to enzyme parameters (*V*_*m*_, *K*_*m*_). Parameters have been chosen here to achieve a baseline ratio of 0.5, indicated with a dashed line. When the flux signal is low, flux will be distributed according to this ratio, but the unregulated pathway is favoured when the flux signal is high. (B) In the feedforward recombine motif, two influxes, *j*_*x*_ and *j*_*y*_, combine to produce flux *j*_*z*_. The product of one of the influxes regulates the other, so that when the regulating influx is high, it is the only source of *j*_*z*_. Conversely, when the signal is low, the nonregulating influx is uninhibited and all of *j*_*z*_ can come from this branch. *K*_*I*_ = 0.05, *V*_*Y*_ = *V*_*Z*_ = 20 and *K*_*Y*_ = *K*_*Z*_ = 0.1 here, and all ratios are at steady state.

Flux entering a split node will be distributed among outfluxes according to the kinetic parameters of the downstream enzymes in the absence of regulation (***LaPorte et al., 1984***). If just one of the downstream enzymes is regulated, then the distribution of fluxes among the two branches can be changed as a function of the regulatory signal, which may be a flux. Capacity-modulating enzyme regulation can significantly change the distribution of downstream fluxes, to the point of committing all of the influx to just one pathway. In contrast, sensitivity modulating regulation does not affect the flux distribution as significantly, but does modulate the influx value at which the maximum effect of capacity modulation occurs (see Appendix 2, Fig 1B). We will focus on regulation of enzyme capacity because of this.

In the feedforward split, a flux-sensing metabolite inhibits one branch downstream of the split point at which it is situated so that as flux into the branchpoint changes, the proportion of this flux allocated to either branchpoint changes in response. This system has the benefit of the down-stream flux distribution being coupled to influx before the influx is split. We examined the feedforward split architecture by simulating changing flux entering a split via a substrate of a reversible reaction (*X* in Fig 3A), with the substrate inhibiting one of the branches (flux *j*_*Z*_). We found that the outflux distribution in this structure does indeed reflect upstream flux for a large range of influx values if the reaction consuming *X* is favoured in the reverse direction (*K*_*eq*_ ∼ 10^−2^) and only for extreme influx values when it is favoured in the forward direction (*K*_*eq*_ ∼ 10^3^), consistent with our previous results. Thus if *X* is the substrate of a thermodynamically constrained reaction, its concentration can be used across a wide range of influxes to control flux partitioning at a split; it is a flux sensor and the downstream enzyme actuates the signal it produces.

Only certain ratios of flux partitioning between downstream paths are possible in this architecture. The minimum ratio is that which is defined by the kinetics of the downstream reactions in the absence of regulation (dashed line in Fig 3, see Appendix 2). More flux can be allocated to the unregulated branch, but it is not possible to allocate less than the base ratio when only one branch is regulated. In contrast, in an equivalent feedback configuration, with *Y* or *Z* as the regulatory signal instead of *X*, any flux ratio is possible since the control signal is dependent on downstream effects which are independent of the influx (Appendix 2, Fig 1).

Because this strategy guarantees a certain baseline ratio for resource allocation, it could be useful in the case where both branches produce essential metabolites, but one pathway is more costly, is not always necessary, or operates unstably. When influx is high, indicating resource abundance, it can all be allocated to the production of the more expensive metabolite, but when it is low, the baseline ratio is still guaranteed and minimum requirements can be met. While we have not shown it here, the logic of FFA of one of the branches is similar: a baseline ratio is guaranteed, with flux being drawn off by the activated pathway only when it is in excess. For example, the flux sensing metabolite in this scenario could indicate carbon uptake flux and when this flux is high it could stimulate a larger fraction of flux being drawn off to a storage pathway.

In the recombine architecture, two different fluxes enter at the same node so that the resulting flux comes from some linear combination of them both. The potential for recombine architectures in flux-sensing schemes has been noted previously, but not described in depth (***Okano et al., 2020***). Similarly to the split, we simulated a recombine architecture with one influx branch as an inhibitor of the other, considering both the reverse-favoured and forward-favoured conditions (Fig 3B). This revealed that the full range of flux splits between both influxes is possible, and occurs as a function of the influx of the regulating branch for the reverse-favoured scenario, and for extreme flux values only in the forward-favoured scenario. In this case, the structure has a clear advantage over the equivalent feedback structure in which the product of the outflux branch regulates one of the influxes, because there is no way of knowing the proportion of outflux produced by either branch once it has passed through the recombine node. The feedforward recombine architecture could be useful in cases for which one branch is more expensive than the other, so the latter is preferred. When flux in the preferred branch is high, all of the outflux can come from this branch so that the expensive pathway is only used as necessary. Since the feedforward recombine architecture can achieve any ratio of fluxes, it could also be useful for stoichiometric balancing of substrates of a bimolecular or higher-order reaction.

We have examined only two flux signal interpretation motifs here. These were chosen based on their presence in central carbon metabolism (CCM), as we will show in the next section. However, many more structures are possible and may be capable of integrating flux signals.

### A Few Bottlenecks Reflect Environmental Conditions via Fluxes

To look for evidence of flux sensing structures, we collated metabolomic and fluxomic data sets for *E coli* fed up to 14 different carbon sources and where possible looked directly for flux-concentration relationships by correlating fluxes and metabolite concentrations (see Methods and Materials) (***Gerosa et al., 2015***; ***Kochanowski et al., 2017***; ***Bennett et al., 2009***). We clustered fluxes and metabolites based on these relationships to find that the correlation structure of the network corresponds to the three primary pathways of CCM in *E coli*: glycolysis, the pentose phosphate pathway (PPP), and the TCA cycle and glyoxylate shunt (Fig 4). Each of these pathways has a distinct pattern of correlations between its fluxes and the measured metabolite concentrations across CCM. The correlations within each cluster provide a snapshot of how concentration signals emerge *and* are interpreted. Thus these relationships must be carefully qualified, since they could arise from an upstream flux influencing a concentration as in a flux sensor, mass balance relationships, or regulatory interactions.

**Figure 4.**
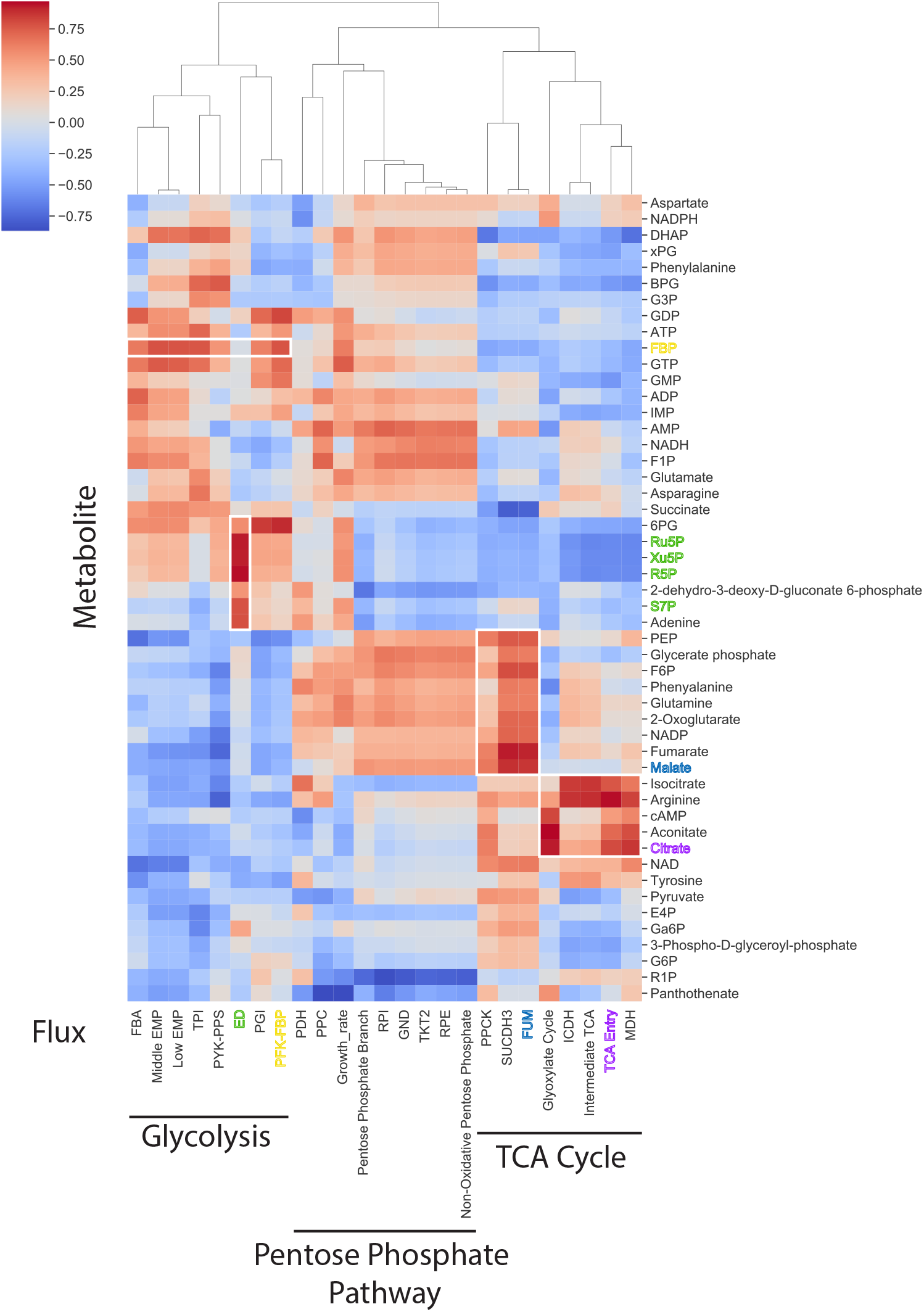
Flux-concentration correlation structure for CCM. Correlations between measured fluxes (bottom) and metabolite concentrations (left) for *E coli* fed eight different carbon sources are presented. Cells indicate the Pearson correlation coefficient for each pairwise interaction, as per the colourbar, and interactions are clustered based on the Euclidean distance metric. Fits are of the form *c*_*i*_ = *a* · *j*_*k*_ + *c*_0,*i*_ for pairs of concentrations (*i*) and fluxes (*k*). Clustering reproduces the three major metabolic pathways in CCM, as indicated. Some fluxes have been lumped and given descriptive names; a full breakdown of the specific reactions in each lumped flux is available in Table 1, Methods and Materials. Colour coding of metabolites and fluxes reflects correlations between metabolites and their producing fluxes, or in the case of the PPP intermediates (green), correlation between concentrations and a related flux (see Fig 5). Relevant local clusters have been highlighted with white boxes. These indicate bottlenecks in which a single constrained flux causes correlations between a few local fluxes and the metabolites they produce.

For example, in the glycolysis cluster, FBP and 6-phosphogluconate (6PG) have the most consistent correlations across fluxes: they are positively correlated with all but one and every glycolytic flux, respectively. However, unlike FBP, 6PG is not correlated with its producing flux (“Pentose Phosphate Branch”, *r* = −0.28), which is clustered separately with the PPP fluxes. 6PG is thus not likely to be a flux sensor. Instead, its concentration is probably influenced by mass balance constraints on glycolytic fluxes. FBP on the other hand has a strong correlation with the phosphofructokinase-fructose bisphosphatase (PFK-FBP) flux (*r* = 0.77, *p* = 0.02). This is consistent with its role as a flux sensor. The fact that it is correlated with all but one of the remaining glycolytic fluxes is due to a combination of mass balance constraints, since these fluxes form a linear pathway, and regulatory interactions since FBP activates pyruvate kinase. FBP is unique in that it is the only metabolite in the glycolysis cluster which is strongly correlated with its producing flux. We posit that this flux-concentration relationship arises because the consuming flux of FBP in the glycolytic mode, FBA, is thermodynamically constrained. It is likely that FBP feeds information forward in the network as a result of its unique position in glycolysis, because it is the only metabolite in the cluster which carries information about upstream fluxes

**Table 1.** Flux short codes and groupings.

| Flux Label | Reaction(s) |
| --- | --- |
| FBA | Fructose biphosphate aldolase |
| Middle EMP | Glyceraldehyde-3-phosphate dehydrogenase, Phosphoglycerate kinase |
| Low EMP | Phosphoglycerate mutase, Enolase |
| TPI | Triose-phosphate isomerase |
| PYK-PPS | Pyruvate kinase, Phosphoenolpyruvate synthase |
| ED | 6-phosphogluconate dehydratase, 2-dehydro-3-deoxy-phosphogluconate aldolase |
| PGI | Glucose-6-phosphate isomerase |
| PFK-FBP | Phosphofructokinase, Fructose biphosphatase |
| PDH | Pyruvate dehydrogenase |
| PPC | Phosphoenolpyruvate carboxylase |
| Pentose Phosphate Branch | Glucose-6-phosphate 1-dehydrogenase, 6-phosphogluconolactonase |
| RPI | Ribose-5-phosphate isomerase |
| GND | Phosphogluconate dehydrogenase |
| TKT2 | Transketolase 2 |
| RPE | Ribulose 5-phosphate 3-epimerase |
| Non-Oxidative Pentose Phosphate | Transketolase 1, Transaldolase |
| PPCK | Phosphoenolpyruvate carboxykinase |
| SUCDH3 | Succinate dehydrogenase |
| FUM | Fumarase |
| Glyoxylate Cycle | Isocitrate lyase, Malate synthase |
| ICDH | Isocitrate dehydrogenase |
| Intermediate TCA | 2-oxoglutarate dehydrogenase, Succinyl-CoA synthetase |
| TCA Entry | Citrate synthase, Aconitase |
| MDH | Malate dehydrogenase |

Source: (Gerosa et al., 2015)

Many strong correlations exist between fluxes and metabolites in the TCA cycle. These form two distinct clusters which are centered around the dehydrogenation of malate catalyzed by malate dehydrogenase (MDH), and the isomerization of citrate catalyzed by the aconitase complex (Fig 4, white boxes) (included here in the “TCA Entry” flux, see Appendix 3). Both malate and citrate correlate strongly with their producing fluxes (*r* = 0.87, *p* = 0.02, and *r* = 0.80, *p* = 0.005 respectively. Fig 5A,B) which is consistent with the fact that MDH (*K*_*eq*_ ∼ 10^−6^) and aconitase (*K*_*eq*_ ∼ 10^−2^) catalyze thermodynamically constrained reactions. These metabolites thus have the potential to be flux sensors for the flux entering and exiting the TCA cycle. Indeed, both metabolites are feedforward regulators (Fig 5C).

**Figure 5.**
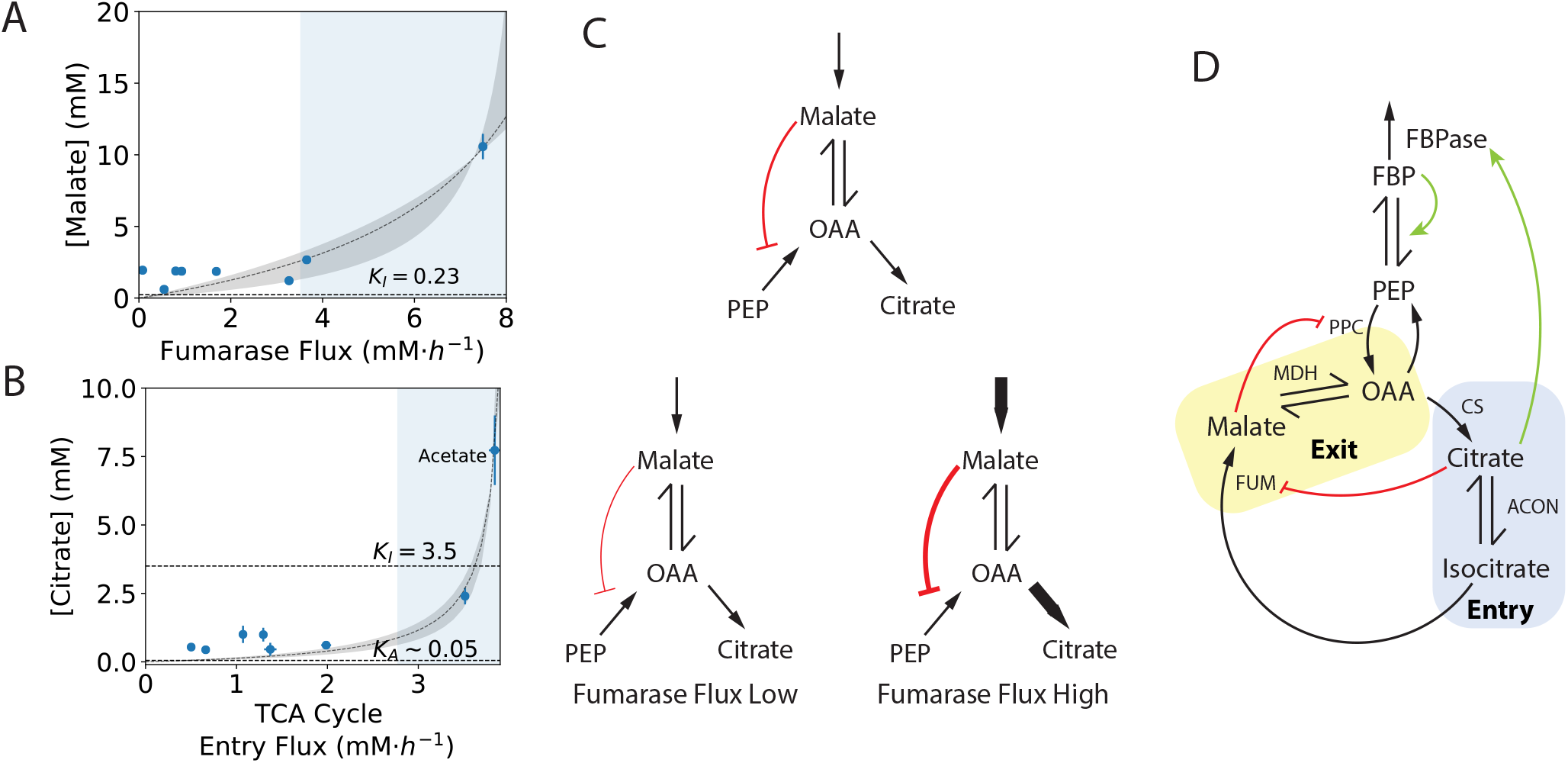
Evidence for novel flux sensors in CCM. Detailed flux-correlation relationship for malate concentration and fumarase (FUM) flux (A), and citrate concentration and TCA cycle entry flux (includes citrate synthase and aconitase, see Table 1, Methods and Materials) (B), with fit to a model of the form 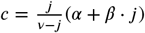 (*R*^2^ = 0.86 for malate and 0.96 for citrate). The grey area represents the 95% CI based on parameter estimates, and the shaded blue region corresponds to the range of fluxes for which the concentration amplifies the flux signal, 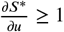. In (A) the inhibition constant for inhibition of phosphoenolpyruvate carboxylase (PPC) is indicated (*K*_*I*_ = 0.23 mM), while in (B) the estimated activation constant for fructose bisphosphatase (FBPase) (*K*_*A*_ = 0.05 mM) and inhibition constant for FUM (*K*_*I*_ = 3.5 mM) are indicated. (C) Malate in a feedforward recombine architecture. OAA may come from malate or PEP. Because malate inhibits PPC, when FUM flux is high malate is prioritized as a source of OAA. (D) Malate and citrate interactions in context. These two bottlenecks naturally break the TCA cycle into two modules which we have called “entry” and “exit”. Flux signals from the entry module are coordinated with flux in the exit module via inhibition by citrate, while anaplerotic flux via PPC is coordinated with exit flux via inhibition by malate. Coordination between modules only occurs to an appreciable degree in cells fed acetate, in which the citrate concentration exceeds the inhibition constant for the citrate-FUM regulatory interaction. Citrate also positively regulates FBPase which could pull flux upward through gluconeogenesis.

Malate inhibits phosphoenolpyruvate carboxylase (PPC) (***Gold and Smith, 1974***) in a feedforward recombine topology (Fig 5B). Inhibition of PPC by malate could prioritize oxaloacetate (OAA) production via the TCA cycle/glyoxylate shunt over the anaplerotic route of PPC to avoid using up PEP except as required (Fig 5C). This interaction is apparently active for all measured conditions in the collated data set since the malate concentration greatly exceeds the *K*_*I*_ value for PPC (0.23 mM) for all conditions (Fig 5A). However, according to our model signal amplification in this case only occurs for fumarase fluxes greater than ∼3 mM·h^−1^, on three of eight carbon sources (Fig 5A, blue region).

Citrate activates fructose bisphosphatase (FBPase) and inhibits FUM (***Hines et al., 2007***; ***Flint, 1994***). It also inhibits phosphofructokinase (PFK), but only the PFKII isozyme (***Kotlarz and Buc, 1981***), which accounts for a very small fraction of the PFK activity in vivo (***Kotlarz et al., 1975***). Activation of FBPase by citrate could act as a feedforward or feedback interaction depending on the direction of carbon flow (Appendix 3, Fig 1). Inhibition of FUM by citrate could coordinate the rate of material entering the cycle with the rate leaving (“entry” and “exit” modules in Fig 5D). The citrate-FUM interaction is an example of a “stacked” interaction, because malate is sensitive to FUM, which is inhibited by citrate, which is in turn sensitive to flux entering the TCA cycle. This stacking could couple oxaloacetate (OAA) source flux prioritization to flux entering the TCA cycle, and it is the first evidence of any kind of structure in the SMR network (SMRN) for *E coli*.

In contrast to the TCA/glyoxylate cluster, there are no strong flux-concentration correlations within the PPP cluster (Fig 4). However, all of the PPP intermediates correlate strongly with Entner-Doudoroff (ED) flux (Fig 6). Of these intermediates, ribulose-5-phosphate (Ru5P), ribose-5-phosphate (R5P), xylulose-5-phosphate (Xu5P), and sedoheptulose-7-phosphate (S7P) both meet the criteria for flux sensors and are feedforward activators of the pyruvate kinase encoded by *pykA* (***Waygood et al., 1974***). However, contrary to our framework the flux they sense is not their direct producing flux, but that of a parallel pathway. We suspect that this is due to the fact that flux entering the 6PG node must either go into the ED or PPP pathway, and is recombined at the glyceraldehyde-3-phosphate (GAP) node which is also an intermediate in the PPP pathway (Fig 6). Thus, PPP intermediates, which are all substrates of thermodynamically constrained reactions, are linked to ED flux via mass balance. Similarly to FBP, in activating pyruvate kinase they could pull flux downward through lower glycolysis when ED flux is high. This case highlights the possibility of flux sensing emerging from more complex structures involving thermodynamic constraints than the simple linear pathway we considered in our explanatory framework.

**Figure 6.**
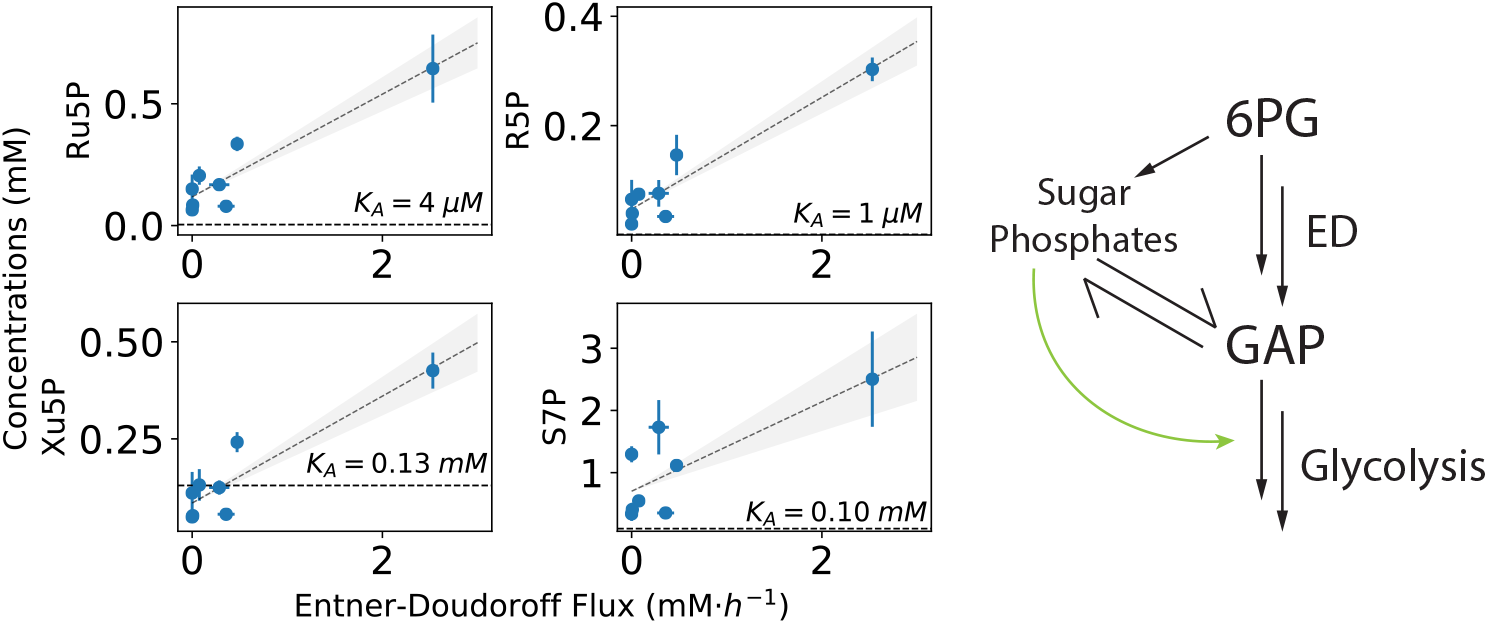
Pentose phosphate pathway (PPP) intermediates as flux sensors. Four PPP intermediates correlate with Entner-Doudoroff (ED) flux and are known activators of the *pykA* pyruvate kinase isozyme (activation constants indicated). Though ED is not a direct producing flux of these intermediates, their fluxes are related by mass balance since all flux entering the 6-phosphogluconate (6PG) node is recombined at the glyceraldehyde-3-phosphate (GAP) node in middle glycolysis whether via the ED or PPP pathways. Each of these flux-concentration relationships operates in the low sensitivity region, 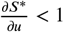 across the collated data set.

We have presented strong evidence consistent with our framework to show that citrate, malate, and PPP intermediates act as flux sensors as a result of thermodynamic constraints. In addition to these examples, we found weak evidence for PEP and glycerol-3-phosphate (G3P) as flux sensors in the collated data. In the case of PEP, variability in the measured phosphoenolpyruvate carboxykinase flux (PPCK) renders the flux-concentration correlation insignificant (*r* = 0.10), while for G3P insufficient natural variation in local fluxes across the carbon sources available prevents a correlation between glycerol uptake and G3P concentration from being made at all. However, G3P satisfies the structural criteria for flux sensing and has a heterogeneous concentration across carbon sources (Appendix 3, Fig 2).

All of the flux sensor candidates we have identified are paired with the entry points to metabolism of just one or two carbon sources. For example, TCA entry-fumarase flux coupling can only occur to any appreciable degree when citrate concentration exceeds 3.5 mM. In the collated data set, this is only the case in cells fed acetate because acetate entry into metabolism is close to the bottleneck point. However, TCA entry-FBPase flux coupling via citrate is apparently relevant for all carbon sources in the collated data set, as is FUM-PPC coupling via malate. This lends credence to our hypothesis that some metabolites – like citrate, malate, and FBP – *always* sense flux because of their position in the network, and this information is used for regulation for specific cases through tuning of regulatory parameters, or differential expression of target enzymes, such as PPC. Other metabolites, such as G3P, may only act as flux sensors in the specific environmental conditions in which their producing flux is non-zero, because this is required for them to accumulate to significantly high concentrations.

## Discussion

As external observers of metabolism, we take for granted how easy our ability to see the whole system renders the performance of even simple analyses of it, such as balancing masses using sensible quantities like concentrations or rates of assimilation of labelled molecules. These quantities can be readily measured and the causes and effects of their observed values can be reconstructed with increasing accuracy and precision, because all of the cellular contents and the environment can be sampled at many time points and sophisticated models with which to interpret this information exist. Cells must perform similar analyses to persist, and they are able to do so on a much shorter timescale and ostensibly much more consistently. Given that amounts are sensible by way of binding, the ability of cells to measure them is intuitive. However, to date, just one limited explanation for how cells are able to measure and respond to changes in rates has been offered (***Kochanowski et al., 2013***).

Building off of the recent discovery and characterization of fructose-1,6-bisphosphate as a key flux-sensing metabolite, we have identified here the physical constraints leading to flux sensing. With our analysis we have shown that constraints on flux capacity are key to the emergence of such flux signals, and thus that without some bottlenecks in the network, flux perturbations would be internally unobservable. This means that the substrates of reactions which are thermodynamically constrained are always sources of flux information to some degree regardless of the regulatory interactions in which they are involved. Context-dependent kinetic constraints on capacity may also yield flux sensing metabolites, but these also enforce infeasibly excessive concentrations, so they are undesirable sources of flux information. In contrast, thermodynamic constraints enforce larger substrate concentrations which are highly sensitive to upstream flux perturbations. Our central hypothesis is thus that thermodynamic bottlenecks in the network yield metabolite concentrations which reflect fluxes, and the information arising from these are valuable signals for the subsequent regulation of other fluxes.

We have demonstrated that this information can be interpreted by real local architectures to make downstream decisions about flux partitioning in the network. The recombine architecture has been mentioned previously, but we have explicitly demonstrated that it can yield arbitrary ratios of influxes as a function of the magnitude of those influxes. We have also shown that the regulation of phosphoenolpyruvate carboxylase by malate is a feedforward recombine structure that may coordinate anaplerosis as a function of TCA/glyoxylate cycle flux. The feedforward split motif we present and characterize is novel, and we have identified two instances in which it may be operational. Beyond these structures, we have also demonstrated that SMR interactions can be stacked, as in the case of fumarase inhibition by citrate. Since malate concentration grows with growing fumarase flux, this stacking could propagate perturbations to TCA cycle entry flux downstream to be sensed as changes in malate concentration. Further work into the dynamics and abundance of stacked regulatory interactions such as this in the small molecule regulatory network could help to elucidate their significance and relevance. Indeed, evaluation of all the exemplar structures we describe, and testing of our hypothesis in general, necessitates targeted experimentation involving deliberate manipulation of rates and quantification of the responses to such manipulations.

We can at best propose our flux sensing theory as a hypothesis here because of the limitations in our approach. We used untargeted data with which it is difficult to separate cause and effect to say with certainty that the flux-concentration correlations we identified arise solely due to thermodynamic constraints. Further, it could be experimentally challenging to verify that these constraints are the specific cause of the flux-concentration relationships we have proposed. However, we find the correspondence between our proposed flux sensors and the known bottlenecks of central carbon metabolism, and their persistence across data sets reflecting natural variation in carbon sources very compelling. Fundamental constraints on metabolism should be consistent across treatments, and this is something we have observed in collating the data sets we used here. Further, the presence of the architectures that we have characterized among the flux sensors we identify is consistent with the framework we have developed. This framework uncovers the potential for the existence of many undiscovered flux-sensing metabolites and flux signal integration structures outside of central carbon metabolism, and for the exploration of the emergence of flux sensing as a universal principle of metabolic regulation. Improved understanding of this principle could enable accelerated engineering of metabolism for a broad range of applications.

## Methods and Materials

### Data Curation and Analysis

We pooled metabolomic data from ***Gerosa et al***. (***2015***), ***Kochanowski et al***. (***2017***), and ***Bennett et al***. (***2009***) by converting concentrations to common units (mM for amounts, mM·h^−1^ for rates) in each data set then taking the grand mean and pooled standard deviation of reported means and standard deviations based on reported sample sizes. We used the conversion factor presented in the Gerosa data set (2.3 mL·gCDW^−1^) for unit conversion of that data. In cases where a metabolite concentration was unavailable in one or more of the data sets we simply only used data from the sets in which it was available and adjusted pooling accordingly. We also excluded data for cells fed glucose + cas amino acids, and those treated with chloramphenicol in the Kochanowski data set for simplicity of analysis and to ensure maximal overlap in the collated data. Since the total concentration of citrate and isocitrate is reported in the Kochanowski data set, we estimated the concentration of either metabolite separately by assuming equilibrium with *K*_*eq*_ = 0.046 (from ***Noor et al***. (***2013***)). This is consistent with the citrate and isocitrate concentrations presented in the Gerosa data set. The final data set is comprised of 51 metabolite concentrations for 14 carbon sources. Of these carbon sources, 3 (glucose, glycerol, acetate) have coverage in all three source data sets, 5 (galactose, succinate, pyruvate, fructose, gluconate) have coverage in the Gerosa and Kochanowski data sets and the remaining 6 (sorbitol, mannose, mannitol, N-acetyl-glucosamine, glucose-6-phosphate, lactate) only have coverage in the Kochanowski data set. Relevant to our analysis is the fact that of these 6 low coverage carbon sources, only one enters metabolism at a point below FBP.

Flux data is only available in the Gerosa data set, so we were able to examine flux-concentration relationships only for flux distributions corresponding to 8 of the 14 carbon sources (first and second categories above). We correlated fluxes and concentrations for each flux-concentration pairing by first assuming a simple linear relationship of the form *c*_*i*_ = *a*·*j*_*k*_+*c*_0,*i*_ with *c*_*i*_ and *j*_*k*_ being concentrations and fluxes, respectively, and *a* and *c*_0_ being parameters. We used least squares regression to evaluate Person’s correlation coefficient for each pairing excluding malic enzyme (ME) fluxes, then clustered pairings based on the Euclidean distance between these coefficients via the Seaborn Clustermap command in Python 3.7. We excluded ME because only four flux measurements are available across carbon sources, and spurious correlations with most metabolite concentrations occurred as a result of this lack of data.

While we expected flux-concentration relationships to be mostly linear, many of the relevant relationships we identified using linear regression ended up being nonlinear. For these, we fit to a model of the form

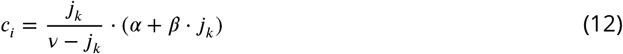

using simple nonlinear regression. Equation 12 is of the same form as equation 6 with *c*_*i*_ being the output concentration, *j*_*k*_ being the input flux for pairings (*k, i*), and *v* being the maximum flux capacity. *α* corresponds with 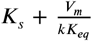 and *β* with 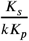. We used these model fit parameters to estimate the lower bound on the sensitive region in Fig 5A, B.

### Dynamic Simulations

Dynamic simulations were used to generate Figs 2 and 3. For both, equations 4 and 5 were used to simulate reversible reactions with parameters specified in figure captions. To avoid infeasible accumulation in the branchpoint system in Fig 2 we used arbitrary saturation kinetics with sufficient flux capacity to remove the products from the system. The kinetic parameters used for this were the same for each branch to facilitate comparison. Where rates or ratios of rates are reported, we first solved the system of ODEs to find steady state concentrations, then used a separate algebraic expression to isolate specific rates of interest

For the systems in Fig 3 we modelled the conversion of *X* to an intermediate using reversible kinetics, and the conversion of that intermediate to products *Y* and/or *Z* using irreversible Michaelis-Menten kinetics. For the recombine architecture, we modelled conversion of *Y* to the intermediate using irreversible kinetics. We balanced kinetic paramaters carefully to ensure that the total flux capacity (i.e. *V*_*m*_) of each step was well above the influxes, *j*_*x*_, provided to the system. Parameters are indicated in the figure caption. Because we found that both capacity-modulating and sensitivity-modulating regulation can achieve a wide range of flux partition ratios (see Appendix), we modelled the regulatory interactions using noncompetitive kinetics, which is a form of purely capacity-modulating regulation.

We used odeint from the Scipy package to solve all the systems presented here, with initial conditions set to zero for all metabolite concentrations and influxes specified as a system parameter. Simulations were performed for sufficient time to achieve steady-state for each influx, and the resulting steady state fluxes were extracted for Fig 3.

### Code and Data Availability

Code is freely available at (future GitHub link) as a Jupyter notebook. Data is available as an additional file at (future eLife link).

## Appendix 1

### The Multiple Consuming Reactions Case

In general, determining the distribution of flux among many branches downstream of a branch point is a complex problem. Flux will distribute according to specific downstream enzyme capacities and sensitivities for influxes well below the capacity of any of the individual downstream paths (***Heijnen et al., 2004***). For our analysis, the exact distribution of flux is not important, because we are primarily interested in how the architecture could accumulate substrate. Thus, we consider the case in which all but one of the consuming enzymes is at saturation. The mass balance is:

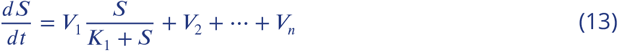

With enzyme 1 being unsaturated and all others operating at *V*_*i*_ = *V*_*m,i*_. This yields a steady state concentration, 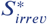, of:

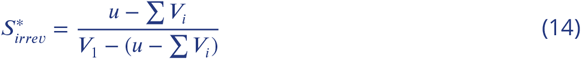

Here the requirement to avoid indefinite accumulation of *S* is *u* < *V*_1_ + Σ *V*_*i*_, so when there are multiple consuming enzymes, feasible steady-state substrate concentrations are guaranteed for a larger range of influxes than if just a single enzyme consumes the substrate. This is intuitive, since infeasbile accumulation in this scenario is dependent on the the total capacity of downstream enzymes to handle influx. Each additional enzyme increases this capacity by *V*_*i*_.

The transformation *u*^′^ = *u* − Σ*V*_*i*_ reveals that the behaviour of 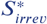 is the same as in the case in which *S* is consumed by a single irreversible enzyme, but over a higher range of influxes, *u* ∈ [Σ*V*_*i*_, Σ*V*_*i*_ + *V*_1_). For fluxes lower than Σ*V*_*i*_, the steady state substrate concentration is guaranteed to be lower than in the case that it is consumed by a single enzyme. As a result, the existence of more paths for consumption also makes kinetically constrained scenarios significantly less likely by shifting the influxes at which they occur to higher values. This shifts the range of reasonable flux signal amplification to higher influx values as well. Now signal amplification occurs for 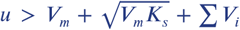. Thus the flux threshold for signal amplification may be significantly larger than in the single enzyme case, and substrates which can be consumed by many enzymes are structurally less likely to be useful signals of flux as a result.

### Region of Infeasible Accumulation

In our model, influx can get arbitrarily close to *V*_m_ with *S*^∗^ and 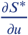 approaching ∞ as it does, so in Fig 1 we arbitrarily chose 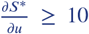 to delineate the range of influxes over which *S* accumulates infeasibly to compare the reversible and irreversible cases. More generally, a reasonable question is for a given choice of α, what fraction of the sensitive region 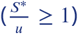 is “lost” to the infeasible accumulation region and is it different between the reversible and irreversible cases?

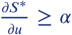 with *α* > 1 occurs for fluxes between 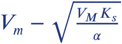 and *V*_*m*_ in the irreversible case, so the total size of this region is:

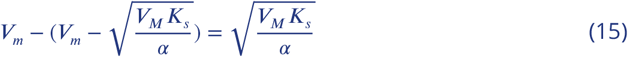

This region corresponds to a fraction of the total amplifying region 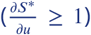, which has a size of 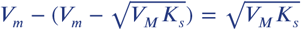. So the fraction of the sensitive region corresponding to infeasible accumulation (defined by a choice of *α*) is:

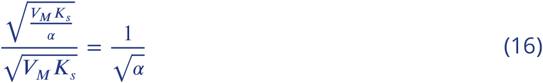

The corresponding fraction in the reversible case (by the same argument) is

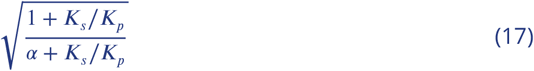

In the irreversible case, this fraction depends only on the choice of *α*, while in the reversible case it is also a function of the parameter group 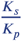. As a result of this, the infeasible accumulation fraction may be larger in the case of reverse-favoured reactions, but in the limit of high relative substrate sensitivity 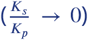, the two scenarios have equivalent fractions. In general, though, for *α* > 1, the reversible case has a slightly larger fraction of infeasible accumulation. Thus, thermodynamic constraints are more likely to yield extreme signal amplification/significant accumulation of substrate than kinetic constraints.

### Dependence of Sensitivity on *K*_*eq*_

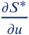 is monotonically increasing for both the reversible and irreversible cases. Therefore

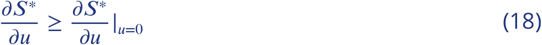

As per equation 11

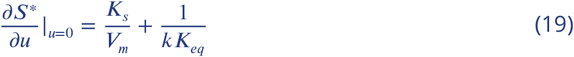

The 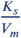 term may take on any non-negative values, so we examine the values of *k* and *K*_*eq*_ which will guarantee 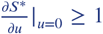, and therefore that the flux signal is amplified for all influx values. If the product *kK*_*eq*_ < 1, then this is guaranteed. This corresponds to

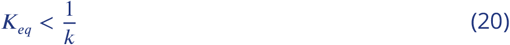

For a downstream turnover of 100 s^−1^, *k* ≈ 100, *K*_*eq*_ must be on the order of 0.01 to ensure that 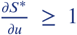. Most enzymes have turnover values significantly lower than this (***Bar-Even et al., 2011***), while there are at least three examples of reactions with *K*_*eq*_ values within this range that we have discussed here (FBA, aconitase, malate dehydrogenase), so there must be at least some examples – and probably many – in metabolism for which favourability in the reverse direction guarantees flux signal amplification.

## Appendix 2

### Splits and Recombines

For the simple case of one flux splitting into two the distribution of downstream fluxes is (***LaPorte et al., 1984***):

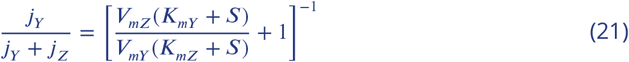

With *S* being the substrate for which downstream enzymes compete, and the *K*_*m,i*_ and *V*_*m,i*_ being kinetic parameters for either competing enzyme, with indices *Y* and *Z* as in Fig 3. This ratio can be controlled by inhibition of just one branch:

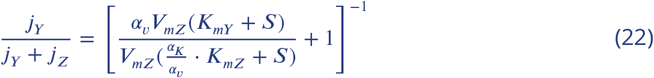

The *α*_*i*_ here are terms characterizing the strength of inhibition by inhibitor 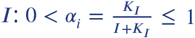, where the *K*_*I*_ are the inhibition parameters unique to each mode of regulation. *α*_*v*_ characterizes modulation of the total capacity of the enzyme, while *α*_*K*_ captures modulation of its sensitivity to substrate concentration. These modes of regulation arise from the ordering of binding of the inhibitor relative to binding of the substrate (***Robin et al., 2018***).

Directly varying *α*_*v*_ reveals that nearly all possible flux partitioning ratios are achievable in the split architecture for some influxes (Fig 1). Varying *α*_*K*_ has a similar effect, but yields a smaller range of flux ratios. Varying either *α* is equivalent to assuming that the effector concentration is not coupled to the influx and that it can vary independently. It is therefore a model of the equivalent *feedback* split, with either *Y* or *Z* as the effector, because the concentrations of these metabolites are not necessarily reflective of upstream fluxes, but instead may be due to other local effects (e.g. downstream fluxes, external uptake, etc), so they can vary independently.

In contrast, when the effector concentration is coupled to the influx in the feedforward configuration – for example, when effector *X* is the substrate of a reverse-favoured reaction – the flux partitioning ratio is also coupled to the flux being partitioned between downstream paths. As a result, there is a baseline flux partitioning ratio, and increases to influx can only cause *more* flux than this baseline to be committed to the unregulated branch. This constrains the possible flux ratios in the feedforward configuration.

That is,

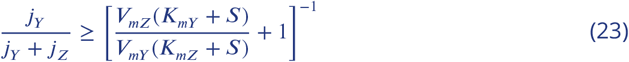

with equality only for very low influx, or very weak inhibition (*K*_*I*_ → ∞). Equation 23 characterizes the baseline ratio for the split motif, which is indicated in Fig 3A as a dashed line. More flux may be committed to the unregulated branch (*j*_*y*_) than the amount determined by the kinetics of the enzyme catalyzing either branch, but a minimum fraction of the total influx is always guaranteed.

**Appendix 2 Figure 1.**
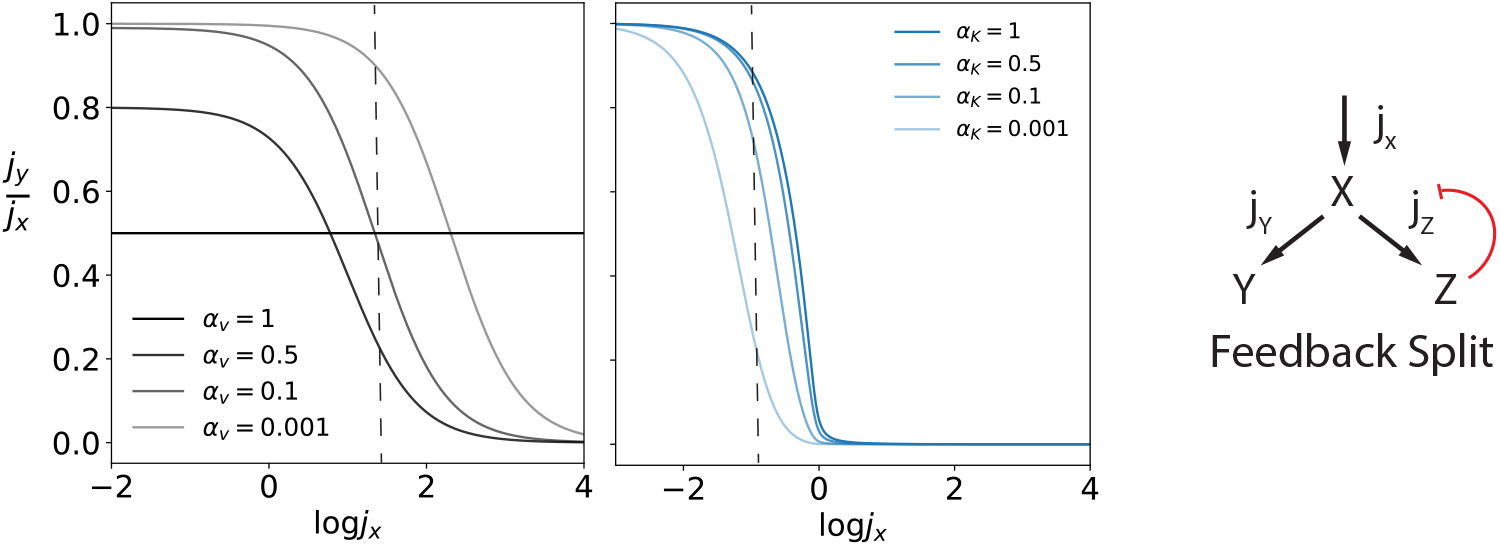
The split motif in the feedback mode. The distribution of fluxes was calculated for various influxes using equation 22 with arbitrary changes to *α*_*v*_ and *α*_*K*_ since the concentration of *Z* is not necessarily coupled to upstream flux, *j*_*x*_. In the absence of capacity-modulating regulation (*α*_*v*_ = 1), the flux distribution does not change with changing influx. However, a wide range of distributions is possible for some influxes (examples indicated with dashed lines) for both types of regulation.

## Appendix 3

### Citrate and FBP as Reciprocal Regulators of Glycolysis

In the glycolytic direction, activation of FBPase by TCA flux-sensing citrate reduces forward flux, both by creating a drive in the opposite direction to net flux and by reducing the FBP signal which activates pyruvate kinase, and thus citrate acts as a feedback inhibitor in this mode. In the gluconeogenic direction, it is an example of feedforward activation because it creates a drive in the same direction as net flux. Structurally, this scheme is analogous to the FBP feedforward regulation motif in which a substrate entering a thermodynamically constrained pathway induces forward pull through that pathway as a function of its influx. While glycolysis and gluconeogenesis have been previously described as reciprocally regulated, the role of citrate as a potential TCA cycle flux sensor highlights how this reciprocal regulation could sense and balance *rates* by measuring amounts at specific points in the pathway.

### PEP and G3P are Potential Flux Sensors

Phosphoenolpyruvate (PEP) is a potential sensor of gluconeogenic flux. Its concentration roughly increases with its producing flux in this metabolic mode, phosphoenolpyruvate carboxykinase (PPCK) (Fig 2A), and it is a feedforward inhibitor of both isocitrate lyase (ICL) and ICDH (***MacKintosh and Nimmo, 1988***; ***Ogawa et al., 2007***), as well as PFKII (***Kotlarz et al., 1975***). The ICL and ICDH interactions are an example of a feedforward split: when PPCK is high, the balance of flux between ICL and ICDH routes could be shifted away from the baseline ratio due to PEP accumulation. The PFK interaction reduces flux in the glycolytic direction, and reduces the FBP signal which activates pyruvate kinase. However, the correlation between PPCK and PEP is not statistically significant in the collated data set (*p* = 0.10) due to variability in the measured value for PPCK flux. Interestingly, we find no evidence that PEP could act as a flux sensor when net flux is in the glycolytic direction. In this case it does not scale with its producing flux, (‘Low EMP’ in Fig 4). Though this may indicate that PEP is a directionally-dependent flux sensor, more work is required to understand if this is indeed the case.

Finally, in the collated data set there is some evidence of glycerol-3-phosphate (G3P) as a sensor of glycerol uptake flux. Its concentration in cells fed glycerol is a high outlier in the G3P concentration distribution (Fig 2B), which is similar to the fact that malate and citrate concentrations are high outliers in cells fed carbon sources which enter metabolism close to these metabolites (succinate and acetate, respectively) (Fig 3), and is consistent with our flux sensing criterion of signal heterogeneity. G3P is also the substrate of a reaction favourable in the reverse direction, and activates FBA in a feedforward split motif. In cells fed glycerol, FBA activity draws flux into upper gluconeogenesis to fuel biosynthetic reactions, so this feedforward split could draw excess glycerol uptake into biosynthetic pathways by modulating the flux ratio above the baseline. However, unlike the other flux sensing metabolites we have identified, observing G3P as a flux sensor in action would require targeted experimentation because it is the earliest bottleneck possible in the glycerol degradation pathway, so there are not many possibilities for natural variation in nearby fluxes as there are for the other proposed flux sensors. Direct modulation of glycerol uptake flux would be required to evaluate G3P function as a flux sensor.

G3P demonstrates the possible dependence of flux-sensing structures on environmental conditions. The flux-sensing activity of FBP is relatively easy to observe because several different carbon sources enter metabolism above FBP at naturally varied rates, and FBP concentration increases with total flux for all of these (***Gerosa et al., 2015***; ***Okano et al., 2020)***.

**Appendix 3 Figure 1.**
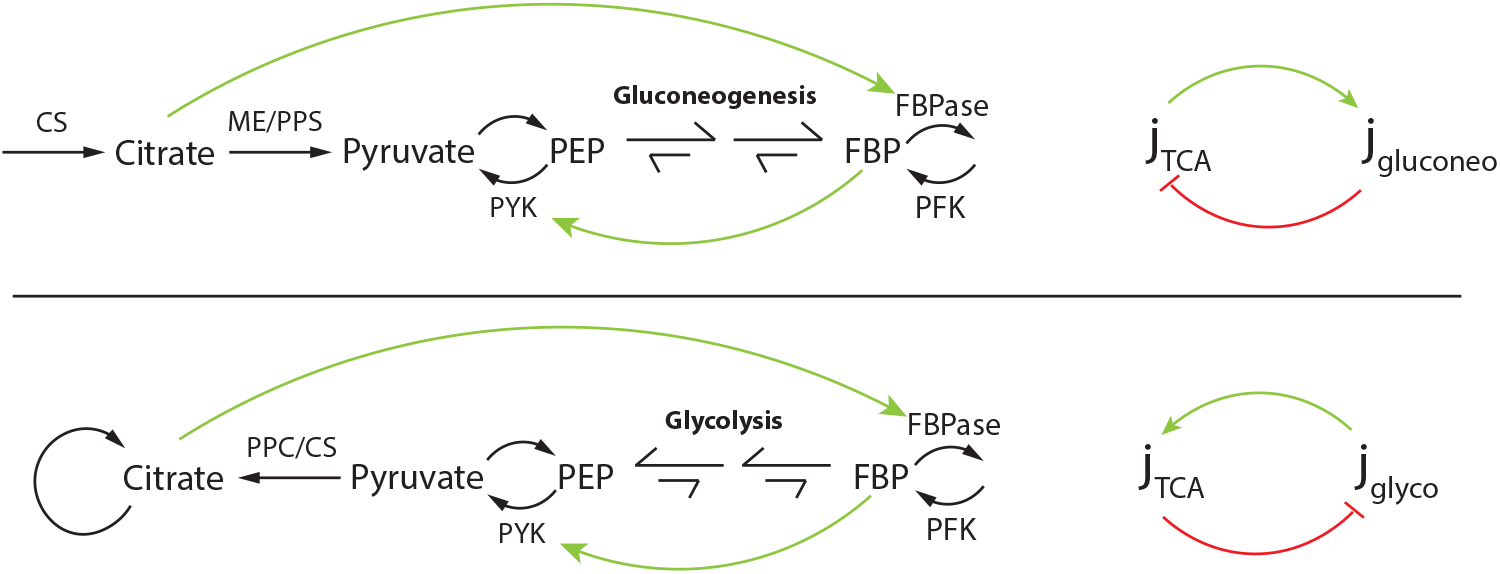
Reciprocal regulation of glycolysis and gluconeogenesis with fluxes as inputs. FBP is a known signal of upper glycolysis flux, and we have shown here that citrate is likely a signal of TCA cycle flux. Citrate is a potent activator of fructose bisphosphatase (FBPase), so it inhibits flux in the glycolytic direction, while FBP activates flux in the glycolytic direction by activating pyruvate kinase. In both the gluconeogenic and glycolytic modes, these interactions yield a feedforward activation-feedback inhibition reciprocal regulatory structure.

**Appendix 3 Figure 2.**
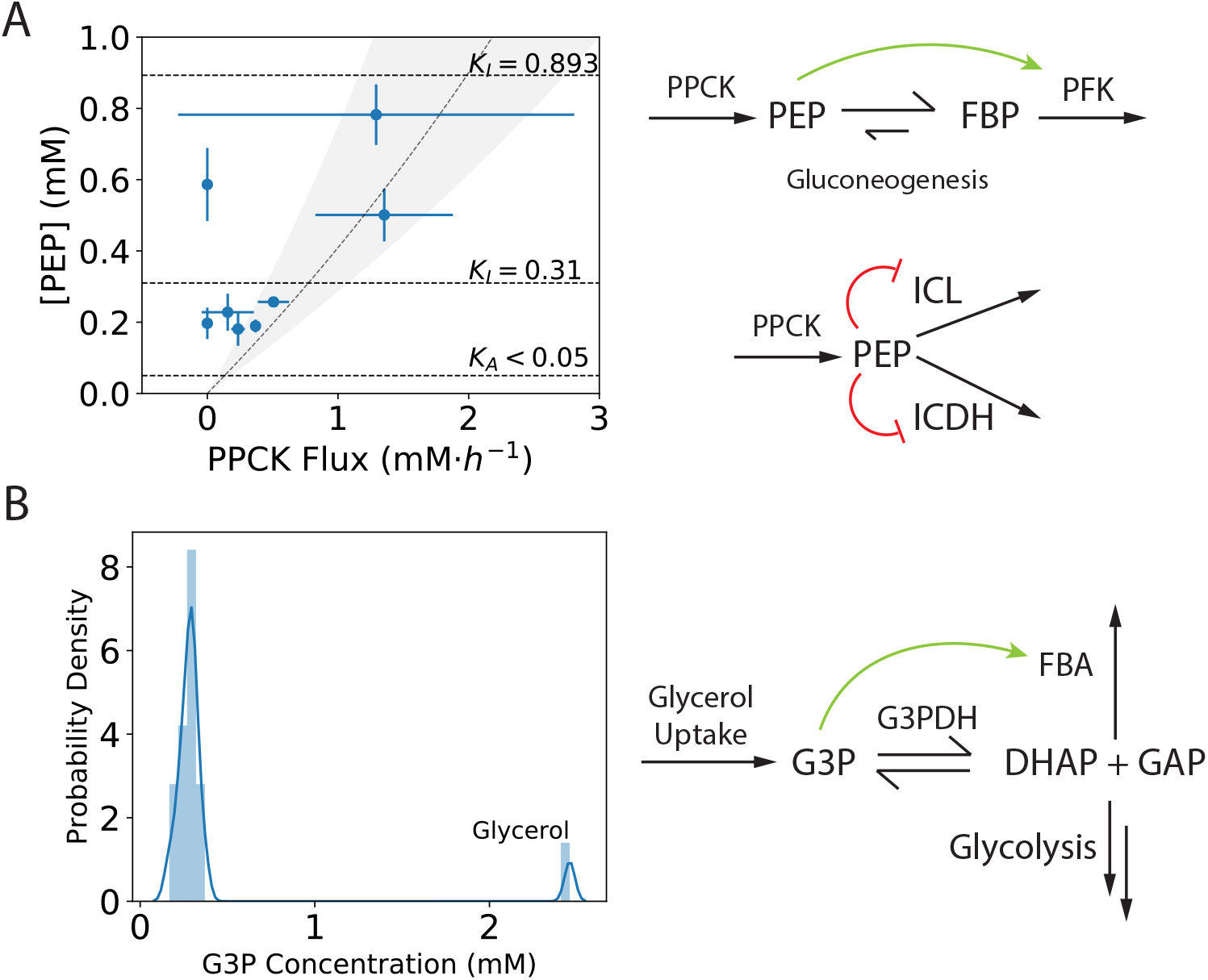
PEP and G3P as flux sensors. (A) PEP concentration grows with its producing flux in the gluconeogenic mode, PPCK, but this correlation is not statistically significant (*p* = 0.10) due to variability in measured PPCK flux at high PEP concentration. PEP is involved in (at least) three feedforward regulatory interactions: it activates phosphofructokinase in a feedforward structure (*K*_*A*_ < 0.5 mM, estimated from data in (***Hines et al., 2007***)), and inhibits isocitrate lyase (ICL) (*K*_*I*_ = 0.893 mM) and ICDH (*K*_*I*_ = 0.31 mM) in a feedforward split architecture. In the collated data set, measured PEP concentration never exceeds the *K*_*I*_ value for ICL. (B) G3P concentration is a high outlier in cells fed glycerol only (Grubbs test, *p* ≤ 0.01). In all other conditions, there is no significant producing flux of G3P, so no obvious correlations emerge in Fig 4. However, the local structure around G3P is consistent with other flux-sensing structures, since the aerobic glycerol-3-phosphate dehydrogenase (G3PDH) has *K*_*eq*_ ∼ 10^−5^. G3P activates FBA in a feedforward split structure.

**Appendix 3 Figure 3.**
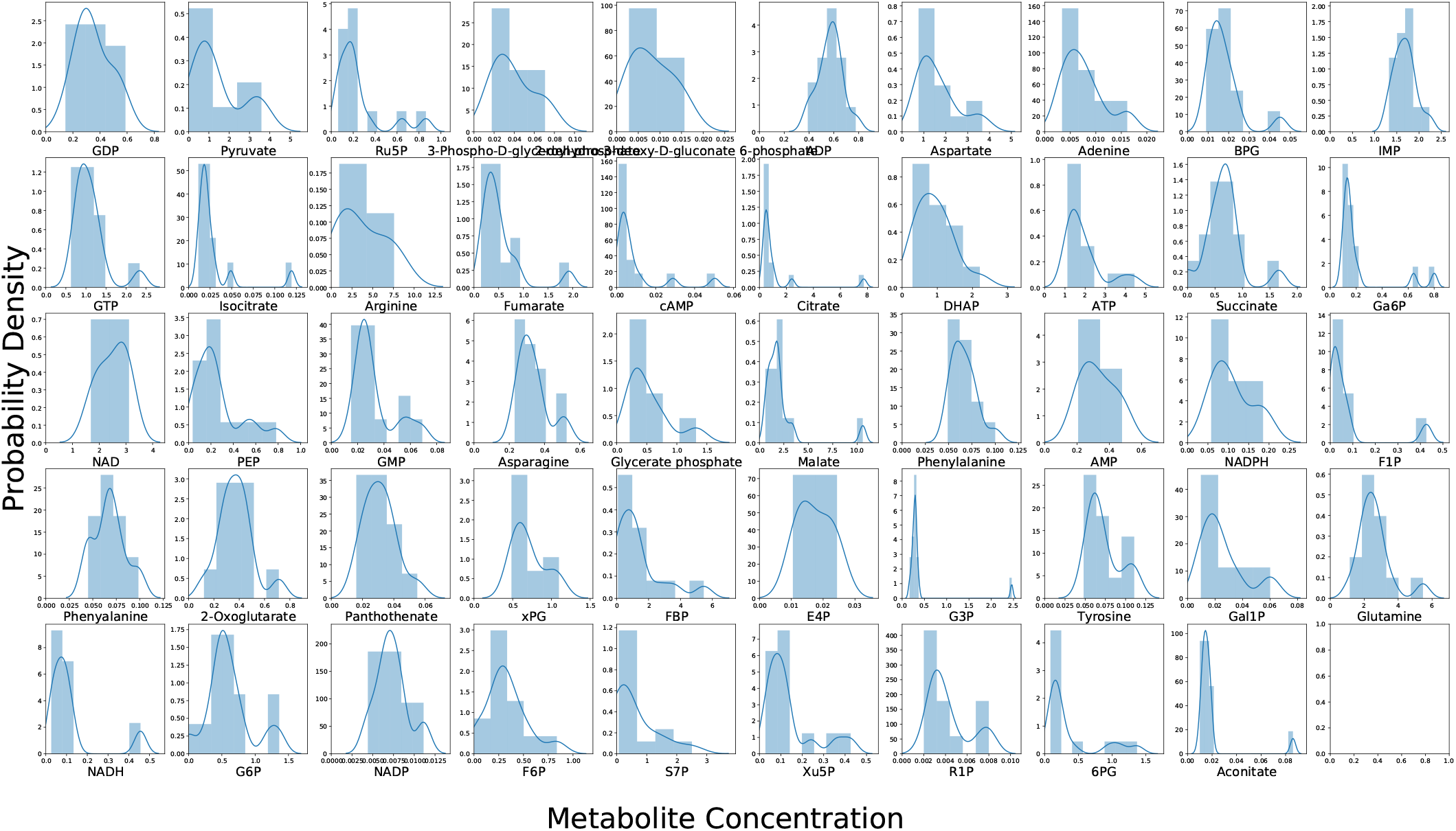
Distributions of metabolite concentrations across conditions in the collated data set. Histograms and kernel density plots are overlaid in each plot. During the exploratory phase we searched for significant outliers using the Grubbs test at a 5% significance level. This revealed that just over half of the metabolites had outlying concentrations in their distributions. Testing at a 1% significance level narrowed this list, yielding citrate and malate and metabolites in their clusters in Fig 4, Ru5P, and G3P, among others. This gave us an indication of context-dependence in the concentration space, and an indication of where to look for flux sensing structures in the network. This variability analysis served as a consistency check with respect to previously published collated data sets. Our list of outlying metabolites at both significance levels overlaps well with those reported in ***Litsios et al. (2018)***.

## Notes

https://github.com/LMSE/flux_sensing

### Competing Interest Statement

The authors have declared no competing interest.

